# A detailed mathematical theory of thalamic and cortical microcircuits based on inference in a generative vision model

**DOI:** 10.1101/2020.09.09.290601

**Authors:** Dileep George, Miguel Lázaro-Gredilla, Wolfgang Lehrach, Antoine Dedieu, Guangyao Zhou

## Abstract

Understanding the information processing roles of cortical circuits is an outstanding problem in neuroscience and artificial intelligence. Theory-driven efforts will be required to tease apart the functional logic of cortical circuits from the vast amounts of experimental data on cortical connectivity and physiology. Although the theoretical setting of Bayesian inference has been suggested as a framework for understanding cortical computation, making precise and falsifiable biological mappings need models that tackle the challenge of real world tasks. Based on a recent generative model, Recursive Cortical Networks, that demonstrated excellent performance on visual task benchmarks, we derive a family of anatomically instantiated and functional cortical circuit models. Efficient inference and generalization guided the representational choices in the original computational model. The cortical circuit model is derived by systematically comparing the computational requirements of this model with known anatomical constraints. The derived model suggests precise functional roles for the feed-forward, feedback, and lateral connections observed in different laminae and columns, assigns a computational role for the path through the thalamus, predicts the interactions between blobs and inter-blobs, and offers an algorithmic explanation for the innate inter-laminar connectivity between clonal neurons within a cortical column. The model also explains several visual phenomena, including the subjective contour effect, and neon-color spreading effect, with circuit-level precision. Our work paves a new path forward in understanding the logic of cortical and thalamic circuits.

## 1. Introduction

Understanding the functional logic of cortical microcircuits is an unsolved problem in neuroscience [1]. In pursuit of this holy grail, advanced imaging and recording techniques are deployed to collect vast amounts of data from the brain [2,3]. Deriving functional cortical circuits from this data is a formidable task, but required to understand how the brain works. To make substantial progress, cortical models need to satisfy three criteria simultaneously [4]: They need to be computationally principled in addressing what the brain does, they need to have algorithmic realizations that solve problems that the brain is able to solve, and they need to have biological implementations that are well grounded in neuroanatomy and physiology such that they explain cognitive and neuroscience observations.

Existing cortical theories and simulations fall short. Large-scale brain simulations [5] aspire to replicate biological details, but lack computational and algorithmic grounding and struggle with solving real-world problems. While the theory of Bayesian inference [6,7] is gaining acceptance as an overarching framework to explain cortical computations, current incarnations primarily focus on describing “what” the brain does, as opposed to “how” it does it [8]. Another class of mechanistic models [9] have the advantage of biological realism, but lack an overall framework and lag in real-world tasks. Overcoming these shortcomings require models that systematically investigate the algorithmic underpinnings of the wide array of questions experimental neuroscientists have answered regarding the connectivity, organization, and physiology of cortical circuits. Each of these experiments provide very important, but partial, clues regarding the functioning of the cortex. Coherent algorithmic models can exploit these cues for useful inductive biases, and help fill in missing pieces, and resolve conflicting information (Fig 1 C).

**Figure 1:**
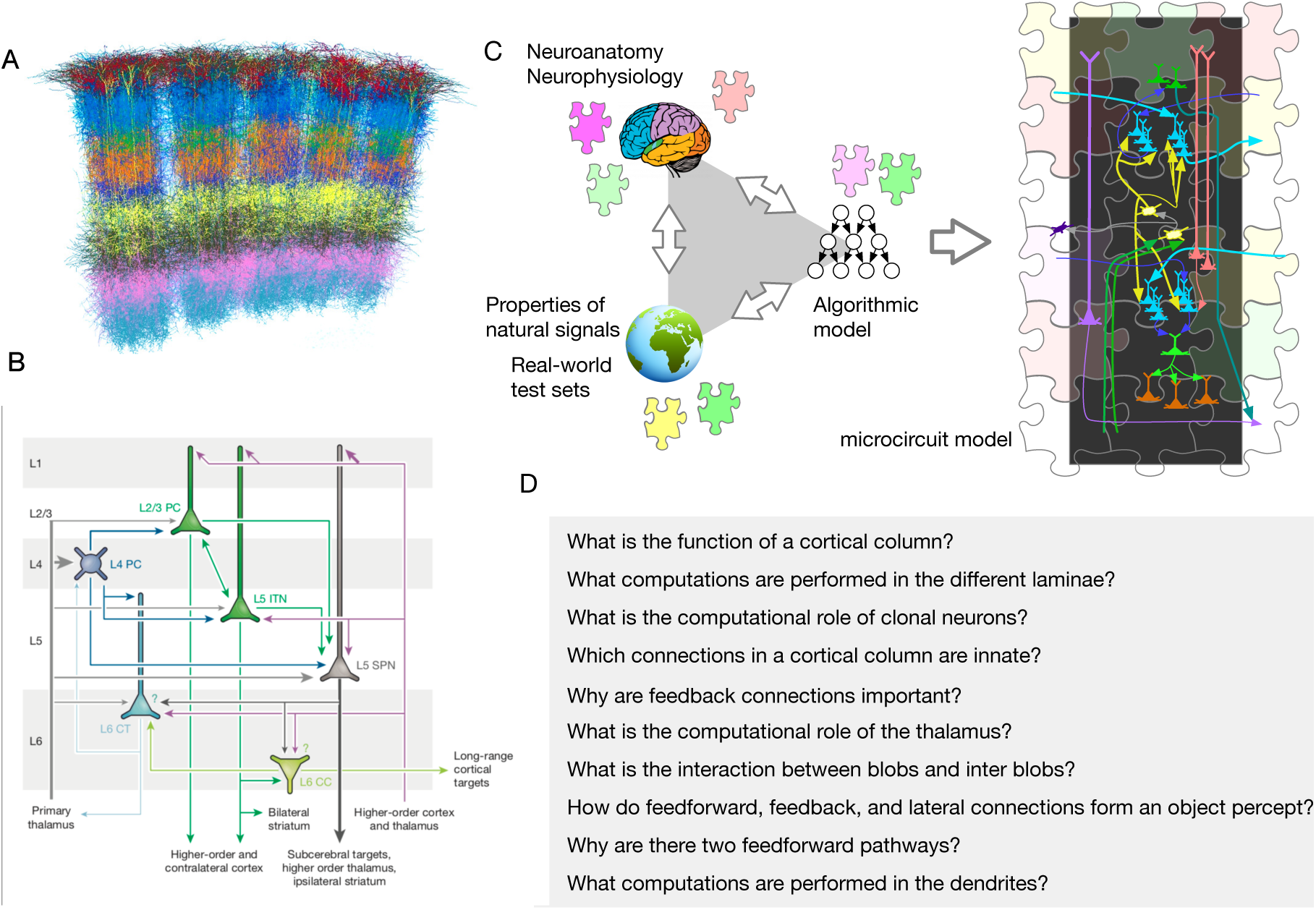
Different approaches to decipher cortical microcircuits. A) A visualization of reconstructed cortical neurons show the formidable complexity in understanding their functional logic. (Image credit [12]). B) A diagram that summarizes cortical connections (reproduced with permission from [13]). The summary is a useful sketch that does not correspond to a functional model. C) Illustration of the triangulation methodology followed in this paper. The brain, the world, and computer science provide partial clues that can be used to construct an overall algorithmic model. The algorithmic model, along with neuroscience data, is then used to derive a functional circuit model. Just like the different pieces in a jigsaw puzzle constrain each other, the combination of algorithmic model and neuroscience data serve to constrain the space of microcircuit instantiations. D) Examples of questions that can be asked and answered with a functional mathematical theory of cortical circuits. This paper answers those questions, and more.

Here, we derive a detailed and functional model of cortical microcircuits based on message-passing inference in Recursive Cortical Networks (RCN), a generative model for vision [10]. RCN is a neuroscience-based probabilistic graphical model (PGM) that systematically incorporated neuroscience findings as inductive biases to achieve state of the art results on several vision benchmarks with greater data-efficiency compared to prevalent deep neural networks.

RCN is consistent with the overarching ideas of Bayesian inference, predictive coding, and free energy minimization [7], but in contrast to prior works that relied on simplistic models [11], the inference algorithms and representational choices of RCN are validated with real-world tasks [10]. High-level Bayesian inference frameworks that do not confront the problem of tractability in realistic settings run the risk of being overly general and unfalsifiable [8], whereas testing on real world settings enable the discovery of architectural and algorithmic details that matter. This distinction is crucial because the inductive biases and representational choices that enable efficient learning and inference also affect the wiring of cortical circuits.

Mapping RCN to a biological implementation results in a detailed model that delineates the role of innateness, learning, and dynamic inference computations in the connectivity and organization of cortical circuits. It assigns computational roles for connections in different laminae and columns, makes precise predictions about the interaction between inter-blobs and blobs, and assigns a functional role to the pathway through the thalamus. The model can also be used to understand the rationale for variations that arise from the same hodology [14]. In addition, the model explains different visual phenomena like subjective contour percept [15,16], and neon-color spreading [17], predicting neurophysiological activations and timing with circuit-level precision. Notably, all these observations are explained simultaneously as the natural byproduct of doing ‘inference to best explanation’ in a model that was learned for parsing a visual scene.

## 2. Recursive Cortical Network (RCN)

RCN is a structured probabilistic graphical model (PGM) for vision consisting of a contour hierarchy of features that interacts with a surface appearance canvas (Fig 2A). The contour hierarchy is learned as alternating layers of feature detectors, pools and lateral connections (Fig 2B). In Fig 2B, each circular node is a binary random variable, the elongated ellipses are categorical random variables, and the rectangles are factors that encode compatibility. Pooling provides invariance to local deformations, similar to the pooling in neocognitron, HMAX model, and convolutional neural nets. The lateral connections between the pools —gray square “factor nodes” in Fig 2 B&C— are learned to enforce contour consistency between the choices in adjacent pools. Pooling provides invariance, whereas lateral connections provide specificity. Fig 2C shows the hierarchical decomposition of a rectangle in terms of simple line segments at the bottom to more complex corner features at intermediate levels. Fig 2D is the graph corresponding to the representation of an “A” from a trained RCN. The graphs corresponding to higher level features share many of their lower level parts as shown in Fig 2B (blue and black), so that a hierarchy of objects is constructed out of many shared parts. The surface appearance canvas, implemented as a Markov Random Field (MRF) as shown in Fig 2E, encodes constraints regarding surface smoothness such as that they are expected to vary smoothly when not interrupted by a contour (nodes in the upper layer), and discontinuously otherwise. A hierarchy consisting of convolutionally tiled graphs of different objects is a probabilistic model for the different objects in a scene.

**Figure 2:**
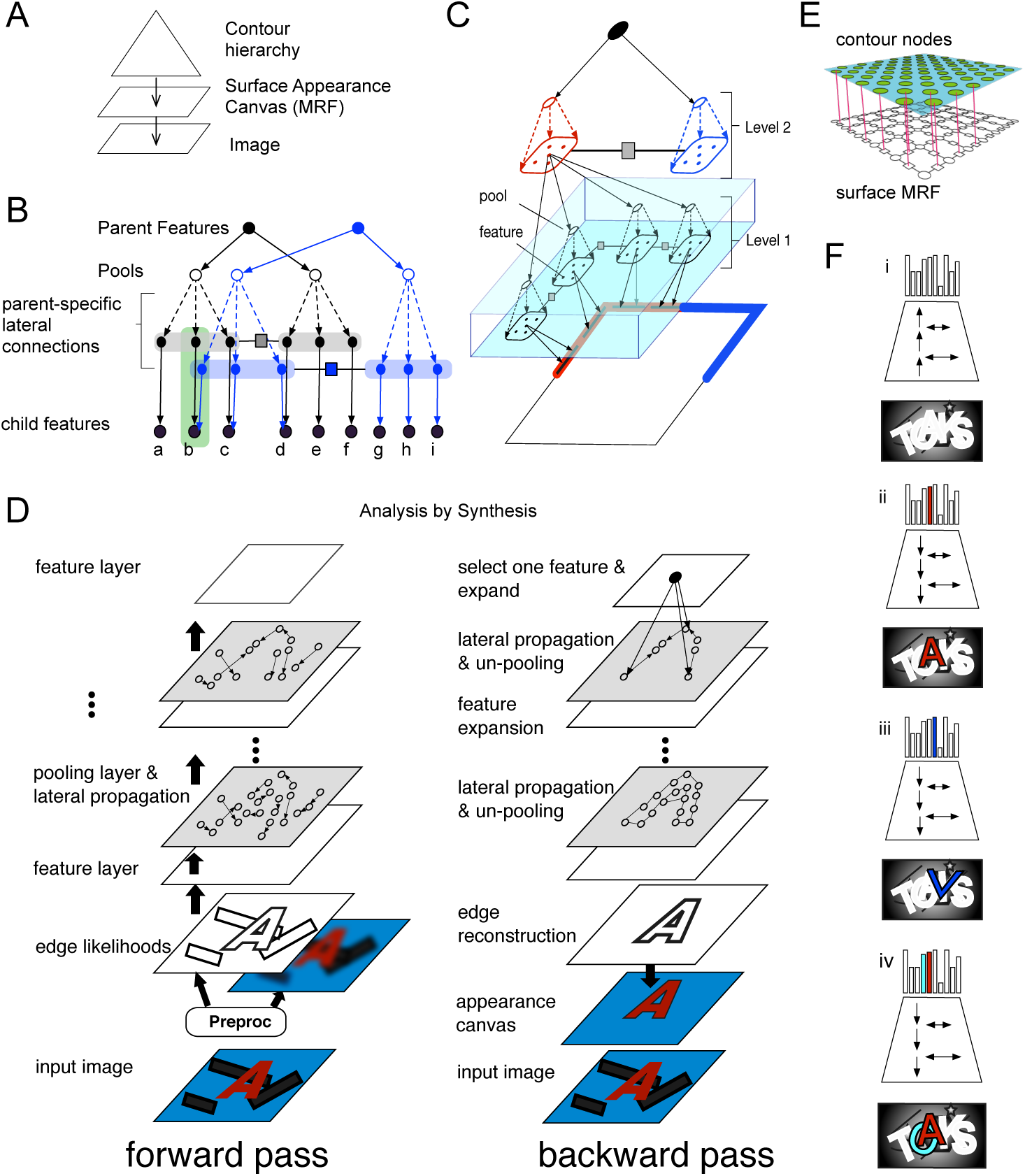
Structure of RCN. (A) A compositional hierarchy generates the contours of an object, and an MRF generates its surface appearance. (B) Contour hierarchy consists of features, pools and laterals. Two subnetworks at the same level of the contour hierarchy keep separate lateral connections by making parent-specific copies of child features and connecting them with parent-specific laterals; nodes within the green rectangle are copies of the feature marked ‘b’. (C) A three-level RCN representing the contours of a square. Features at level 2 represent the four corners, and each corner is represented as a conjunction of four line-segment features. (D) Inference is achieved by passing messages along forward, backward, and lateral directions. (E) The surface appearance MRF. (F) Forward pass identifies object hypotheses in the scene, and backward and lateral pass segments it from the background for analysis by synthesis. Local hallucinations, eg. the ‘v’ in (iii), are explained away during parsing to obtain a global solution that best explains the evidence.

### Message passing inference in RCN

RCN uses belief propagation (BP) [18], a local message passing algorithm for answering global inference queries in a graphical model. Although BP has no theoretical guarantees for producing the right solution in loopy graphs like RCN, appropriate scheduling and damping of messages have resulted in revolutionary empirical success [19].

Scene parsing in RCN is achieved through approximate MAP inference (inference to best explanation) using the max-product [18] version of BP with a schedule inspired by biology (Fig 2 D&F). A fast forward pass, which includes short-range lateral propagations, identifies nodes that are highly likely given the evidence. The backward pass focuses on highly active top-level nodes and includes longer range lateral propagations. Analysis by synthesis [20] is naturally achieved through distributed message passing. The forward and backward passes assemble an approximate MAP solution that produces a complete segmentation of the input scene. See [10] for more details.

### Neuronal implementation of message passing inference

Understanding the neuronal implementation of message-passing inference in a simple graphical model will serve to understand how RCN computations are mapped to biological implementation. Fig 3A shows a simple graphical model with three feature nodes *a, b, c* and three pixel nodes *f, e, g*. When a feature is ON, the pixels it is connected to are ON. When a pixel node has multiple feature nodes connecting it, the interact through an OR mechanism as shown in the corresponding factor graph in Fig 3B. Messages pass along the edges in both directions. Fig 3 C&D show the propagation dynamics of explaining away between the feature nodes when all the pixels are ON. The first forward pass (Fig 3 C) broadcasts the evidence to each parent. Since feature *b* being on explains all the features being on, the backward pass and subsequent forward pass produces explaining away through the computations in the OR factors to diminish the evidence flowing to the nodes *a* and *c*. The explaining away computations in the OR factor *d* are shown in Fig 3F.

**Figure 3:**
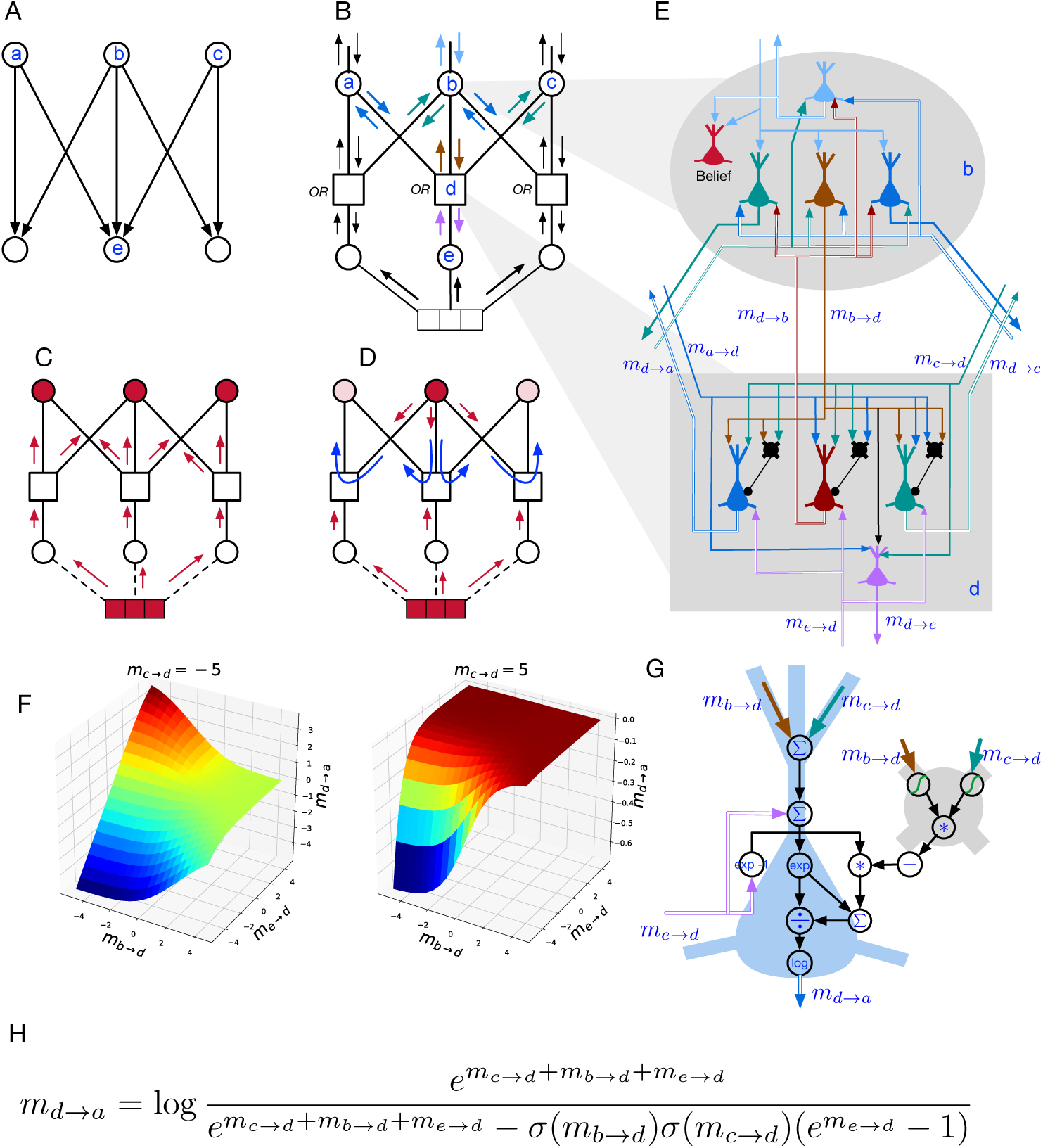
Message-passing and its neuronal mapping in a simple graphical model A) A bipartite PGM with parent nodes representing features, and child nodes representing pixels. The multi-parent interactions at child nodes are modeled by the *OR* function. B-D) Factor graph representation of the PGM in A, with different message annotations. Corresponding to each edge there are two BP messages, one in each direction. Nodes consume the incoming messages to produce output messages. C) First Forward pass copies the bottom-up evidence to parent nodes. D) Subsequent backward-forward passes produce explaining-away competition between parent features. E) Neural implementation of message-passing in nodes *b* and *e*. F) *m*_*d→a*_ as a function of *m*_*e→d*_ and *m*_*b→d*_ shows the excitatory-inhibitory interactions between the inputs. G) Details of computation within an excitatory-inhibitory pair of neurons in *d*, to calculate *m*_*d→a*_, based on the BP equation in H.

Figs 3 E&G show details of how the equations in factor node *d* (Fig 3H) are implemented using neurons. Since neurons are unidirectional, multiple neurons are required within each variable/factor node to convert the incoming messages to outgoing messages. In general, the message to each neighbor is implemented using a different neuron, and carried by its axon. Within a factor node, the synapses between incoming axons, and the dendrites of outgoing axons and interneurons implement the computations required for the factor. In node *d*, the message to each parent is computed by a different neuron, and each of those neurons receive inputs from other parents. The computation involves excitatory and inhibitory interactions between these input connections, as shown in the equation in Fig 3H. Implementing these equations require specific non-linearities and dendritic computations [21] in these neurons, as shown in Fig 3G. The neuronal implementation of distributed probabilistic inference requires precise excitatory-inhibitory interactions and dendritic computations, even for a simple factor graph. Further details and degrees of freedom in such neuronal implementations are discussed in the supplementary material.

## 3. Anatomical Mapping Results

In the following sections we describe how RCN computations (Fig 5) map to cortical circuits, explaining or predicting their functional logic. We will refer to the cortical implementation by the name Bio-RCN, and the abstract graphical model by the name RCN. We first describe the high-level mapping between RCN and Bio-RCN using Figures 4 & 5, and then the details of the microcircuits. Details of the biological implementation are in Figures 6 & 7.

**Figure 4:**
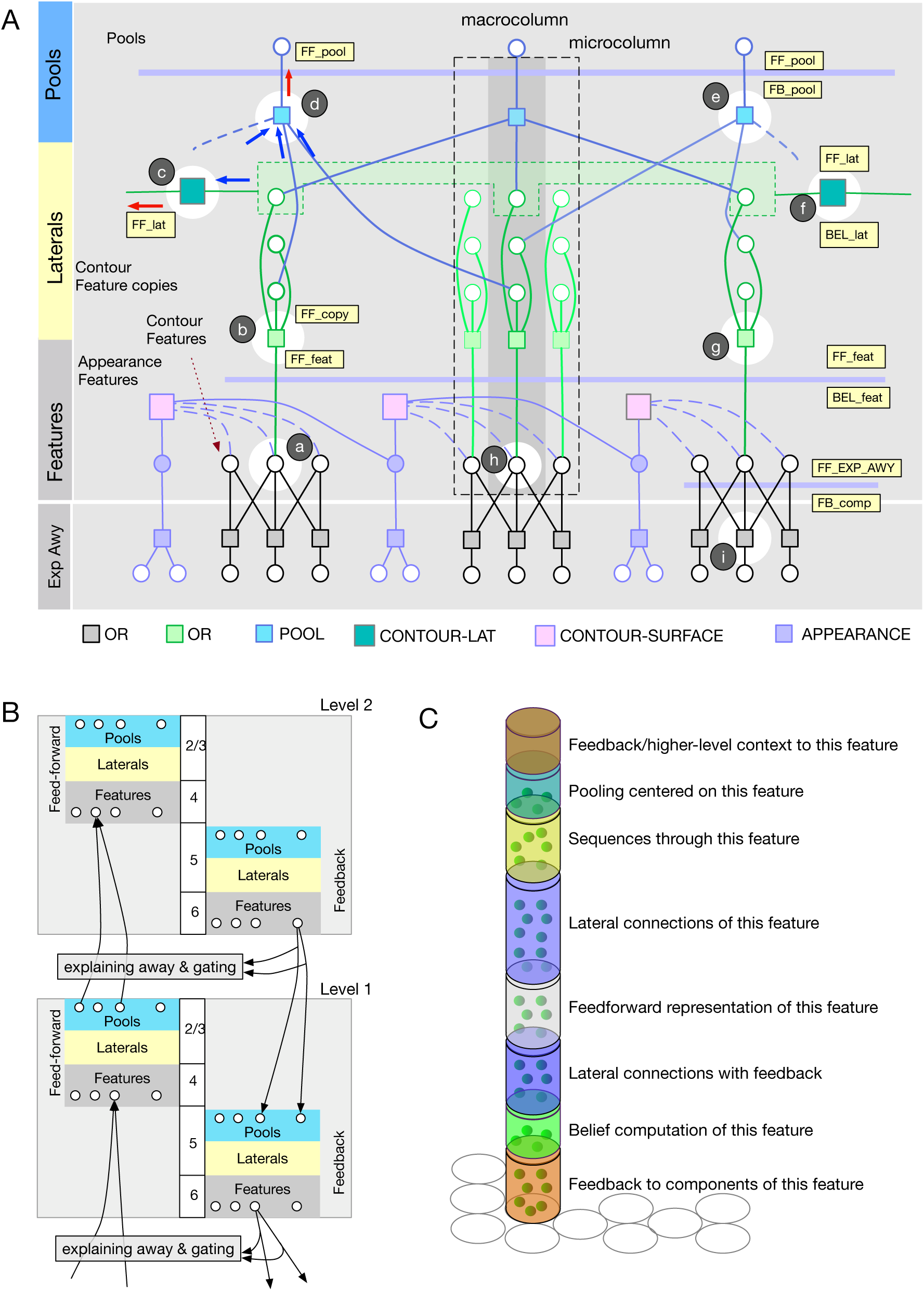
Stages of mapping RCN inference to cortical implementation. (A) Detailed factor graph of a RCN level. Thin horizontal lines indicate messages at different stages of inference, and these map to different laminae in a biological implementation. Variables and messages in a vertical slice as marked map to computations within a cortical column. (B) Conceptual cortical implementation of message passing. Messages in the different directions along the edge of a PGM are implemented in biology using two different sets of neurons for features, laterals, and pools. Messages between different levels go through a stage that includes explaining away and gating. (C) Cortical column as a binary random variable that represents a ‘feature’ or a ‘concept’, for example, an oriented line segment in V1 or the letter ‘B’, in IT. The different laminae in a column correspond to the inference computations that determine the participation of this feature in different contexts: laterally in the context of other features at the same level, hierarchically in the context of parent features, hierarchically as context for child features, and pooling/un-pooling for invariant representations.

**Figure 5:**
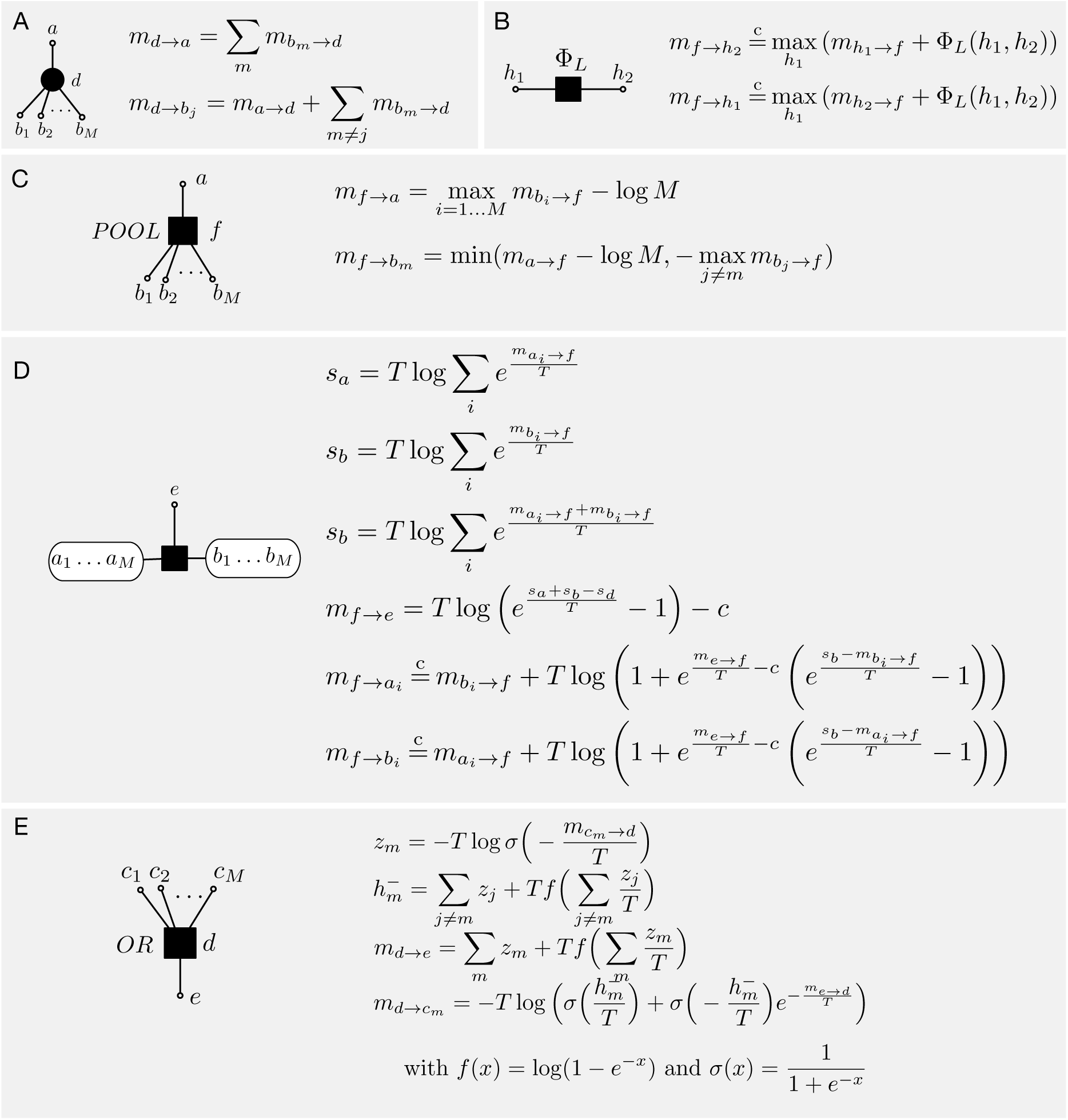
BP message computations in RCN factors and nodes. All computations are in the log domain. Equations with a temperature *T* can do marginalization (*T* = 1), maximization (*T* = 0), or a soft maximization depending on the temperature setting. See supplement for derivations.

**Figure 6:**
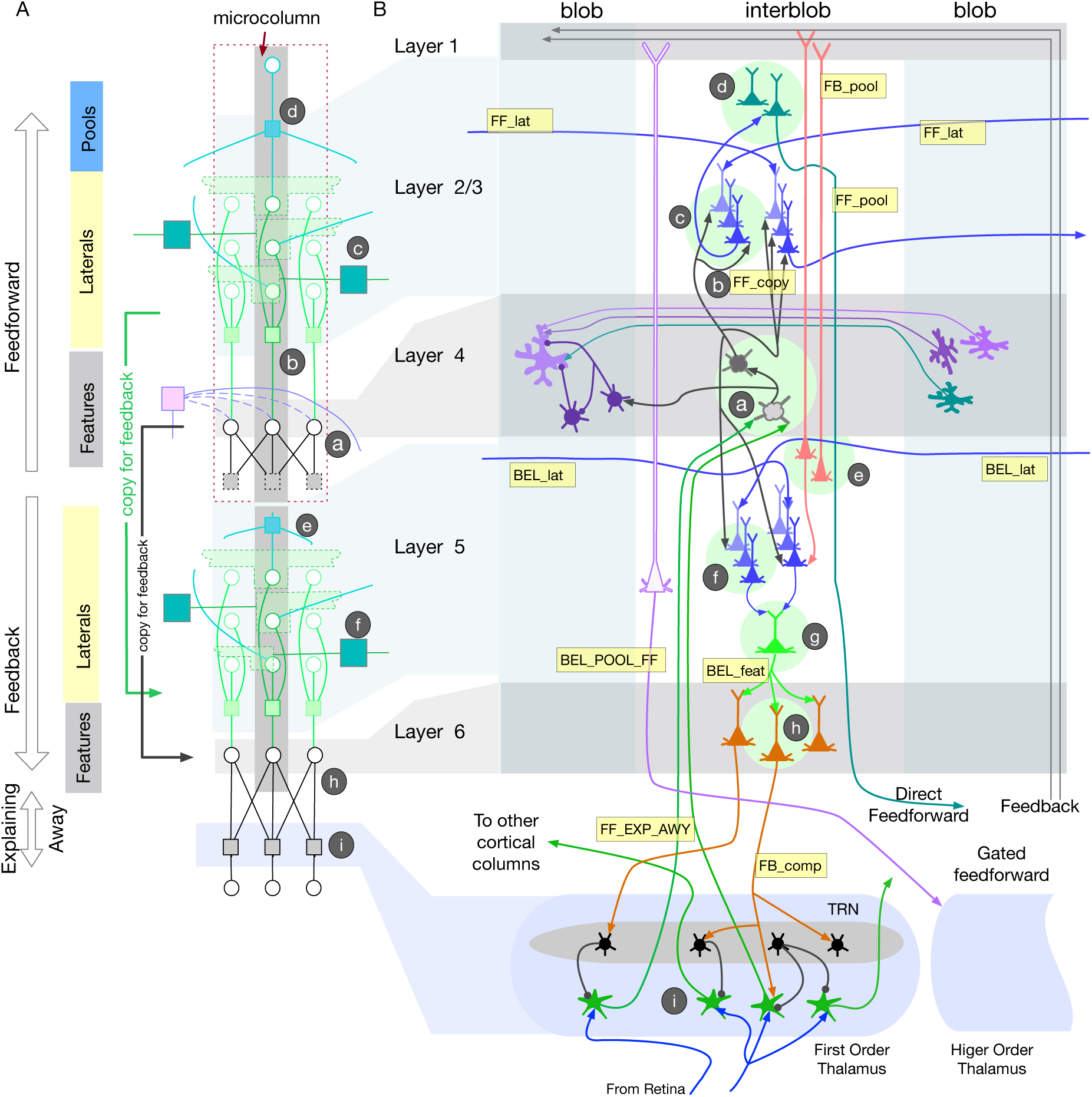
Biological implementation of RCN inference. Top left: RCN factor graph segment for a microcolumn. Since Bio-RCN requires different copies of neurons for forward and backward computations, the factor graph segment is replicated in bottom left to show the correspondence between cortical layers and computations in the factor graph. The correspondence between RCN computations and Bio-RCN are annotated using circled letters 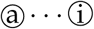

### Overall organization of messages between cortical regions and the thalamus

Fig. 4A shows details of one level of RCN in the form of a factor graph: circles are random variable nodes, and rectangles are factors that encode the dependencies between variables. The variables in one RCN level are contour (black circles) or surface (filled circles) features, contour-copies (green circles) used in lateral connections, and pools (blue circles) that are locally invariant representations of contour segments. A node in the graph consumes incoming messages, processes them and produces outgoing messages on all the edges it is connected to. The variable nodes and the factor nodes implement different computations on these messages, as described in Fig 5. A pool variable of one level in RCN can participate in multiple higher-level features, and their competition is resolved during inference time using computations in the explaining away block.

Fig 4B shows a coarse conceptual diagram of how message passing will be organized in a hierarchy when the feed-forward and feedback messages are implemented on separate pathways in cortical laminae. Within each level (cortical region) neurons in the feed-forward pathway process messages through a features-laterals-pools cascade to send messages to the next level, and the neurons in the feedback pathway do this in reverse, optionally combining feed-forward and feedback at different stages. Cortical laminae 2/3, and 4 typically process feed-forward information exclusively, while laminae 5 and 6 process feedback and produce ‘beliefs’ that combine feed-forward and feedback. In Bio-RCN, these computations will be carried out in the dendrites and soma of networks of neurons. Neuronal outputs at different laminae correspond to messages computed at different stages in the RCN factor graph. Explaining away and gating are implemented in a different block that sits at the interface between two levels.

### Cortical micro-column as a binary random variable

A fundamental organizational unit of neocortex is that of cortical columns formed by synaptically connected vertical clusters of neurons [22], and the computational role of this columnar connectivity has remained a mystery. RCN assigns a functional and computational role to cortical columns: the whole column is viewed as a binary random variable, and the neurons in different laminae perform computations that infer the posterior state of this random variable from available evidence. The random variable represented by a cortical column can correspond to a ‘feature’ or a ‘concept’—for example, an oriented line segment in V1 or the letter ‘B’, in IT. The different laminae in a particular column correspond to the inference computations that determine the participation of this feature in different contexts: (1) laterally in the context of other features at the same level, (2) hierarchically in the context of parent features, (3) hierarchically as context for child features, and (4) pooling/unpooling for invariant representations (Fig 4C). During inference, their activities represent the contributions of these different contextual evidence in support of the feature being ON. The belief that the cortical column is ON itself is represented by specific neurons in specific laminae. In the factor-graph specification of RCN, the different aspects of a feature like lateral membership or pool membership are represented by binary variables themselves, but these binary variables are copies internal to the micro-column, and these copies represent the different contextual interactions of the feature-variable represented by the micro-column.

### Clonally related neurons in a cortical column

RCN assigns a functional rationale for clonally related excitatory neurons derived from a common progenitor. In the ontogenetic column clonal neurons are known to share similar physiological functions, such as visual orientation selectivity [23–25]. RCN uses copies of the same feature to represent its participation in different lateral and hierarchical contexts, and such a representation was shown to be parsimonious and advantageous in representing higher-order spatial or temporal relationships [10,26,27]. The different copies in each group have identical bottom-up inputs or inter-laminar inputs, while they differ in their lateral connections. In RCN, these vertical and top-down connections to the copies of a feature can be established a-priori, without any learning. In Bio-RCN these apriori connections correspond to the vertical connections established in ontogenetic columns (eg. axons marked `FF_copy` in Fig 6) between clonally related neurons. Bio-RCN predicts that several inter-laminar connections within a column can be established apriori as part of development without any learning, and do not need to be modified further with experience, consistent with cortical measurements showing different levels of plasticity for different connections [28]. Recent neuroanatomical observations about clonal neurons [29] are consistent with this proposal. Bio-RCN suggests that a cortical micro-column is initialized by hard-wired developmental programs to have integration of vertical inputs from different copies of the column’s feature, and learned lateral connections from other cortical columns.

### Blobs and interblobs

Contour-surface factorization in RCN maps directly to interblob columns and cytochrome oxidase blobs in the primary visual cortex [30]. The interactions between interblobs and blobs will be considered again in a subsequent section. Unless otherwise specified, descriptions of laminar processing within a column are with respect to interblob columns that represent contour features.

### 3.1. Layer 4: Feed-forward feature evidence computation

According to Bio-RCN, layer 4 stellate cells compute the feed-forward evidence for features (`FF_feat` in Fig 4 A), obtained by the summation of bottom-up input messages `FF_EXP_AWY` along with the messages from contour-surface factors (Fig 5A, annotation 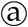 Fig 4 A). Computing this requires direct access to the bottom-up evidence, a requirement matched by spiny stellate cells in layer 4 which are the primary recipients of feed-forward inputs to a cortical region [31,32]. The outputs from layer 4 are intra-columnar projections to layer 2/3 and 5, satisfying the requirements for subsequent computations in RCN.

This mapping assigns precise meaning to the thalamo-cortical axons, and to the different stages of processing in layer 4. Each incoming message, carried by an axon from relay cells in the thalamus (green axons marked `FF_EXP_AWY` in Fig 6B), is a scalar representing the log-likelihood ratio of a component of a feature being on. `FF_feat` is calculated in two stages, shown by gray neurons annotated 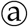 in Fig 6, in Bio-RCN. The first stage computes the likelihood of contour features alone, ignoring the interaction with appearance blobs. The second stage combines this with inputs from the appearance blobs. A multi-stage computation within layer 4 with inputs from blob neurons is consistent with observations [33,34]. See supplement for further evidence.

### 3.2. Layer 2/3: feed-forward lateral connections for contour continuity

Bio-RCN suggests that a subset of pyramidal cells in layer 2/3, annotated 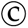 in Fig 6 B, and their inter-columnar lateral projections implement contour continuity inference. Lateral factors in RCN (Fig 4A, annotation 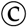) encode a higher-order association field over contour features, and lateral message-passing encourages contour-continuous solutions. In lateral propagation, the likelihood of each feature is calculated as a combination of bottom-up inputs from the features (`FF_feat` and `FF_copy`), and lateral messages (`FF_lat`) from other pools. In Fig 4A, the green colored nodes in the same column are ‘clones’ that receive the same bottom-up input, but have different lateral connections. The clones serve to encode long-range lateral dependencies [35,36], as opposed to just the first order dependencies captured in pairwise models.

Layers 2 and 3 match the anatomical constraints for implementing these computations. Layer 2/3 pyramidal neurons receive feed-forward inputs from the layer-4 neurons [14], lateral inputs from co-circular pyramidal neurons in other columns [37,38], and send their axons across columns covering large distances, making patchy connections at their destinations [39–43]. While the vertical inter-laminar connections column can be specified apriori without learning, the recurrent lateral connections between pyramidal cells are learned from experience [44], consistent with the learning stages in RCN.

The blue neurons annotated 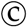 in layer 2/3 of Fig 6 show the biological implementation. All of them receive the same bottom-up input `FF_copy`, axon branches of `FF_feat` calculated by layer 4 neurons. The two sets correspond to messages propagating in two different directions, and the copies within each set help to encode higher-order dependencies [26,27]. Anatomical data supports this proposal: layer-4 spiny neurons make focused and dense connections to layer 2/3 pyramidal cells, with each spiny neuron innervating 300-400 layer-3 pyramidal cells [32].

The lateral factor is a sparse matrix that encodes the compatibility between the features in the different pools. In Bio-RCN this factor is realized in the dendritic trees of the neurons involved, implementing the computations in Fig 5 xx. The lateral factors treat each pool as a categorical random variable that introduces mutual exclusivity between the constituent binary feature random variables (light green region in Fig 4A). This operation involves mutual inhibition and normalization between the features that form constituents of the pool. We suggest that somatostatin-expressing inhibitory neurons (SOMs) in the superficial layers of the cortex are suited for this per-pool normalization, consistent with recent findings of [45]. Fig 7B shows a more detailed implementation of the lateral connections between two pools, including their inhibitory fields.

**Figure 7:**
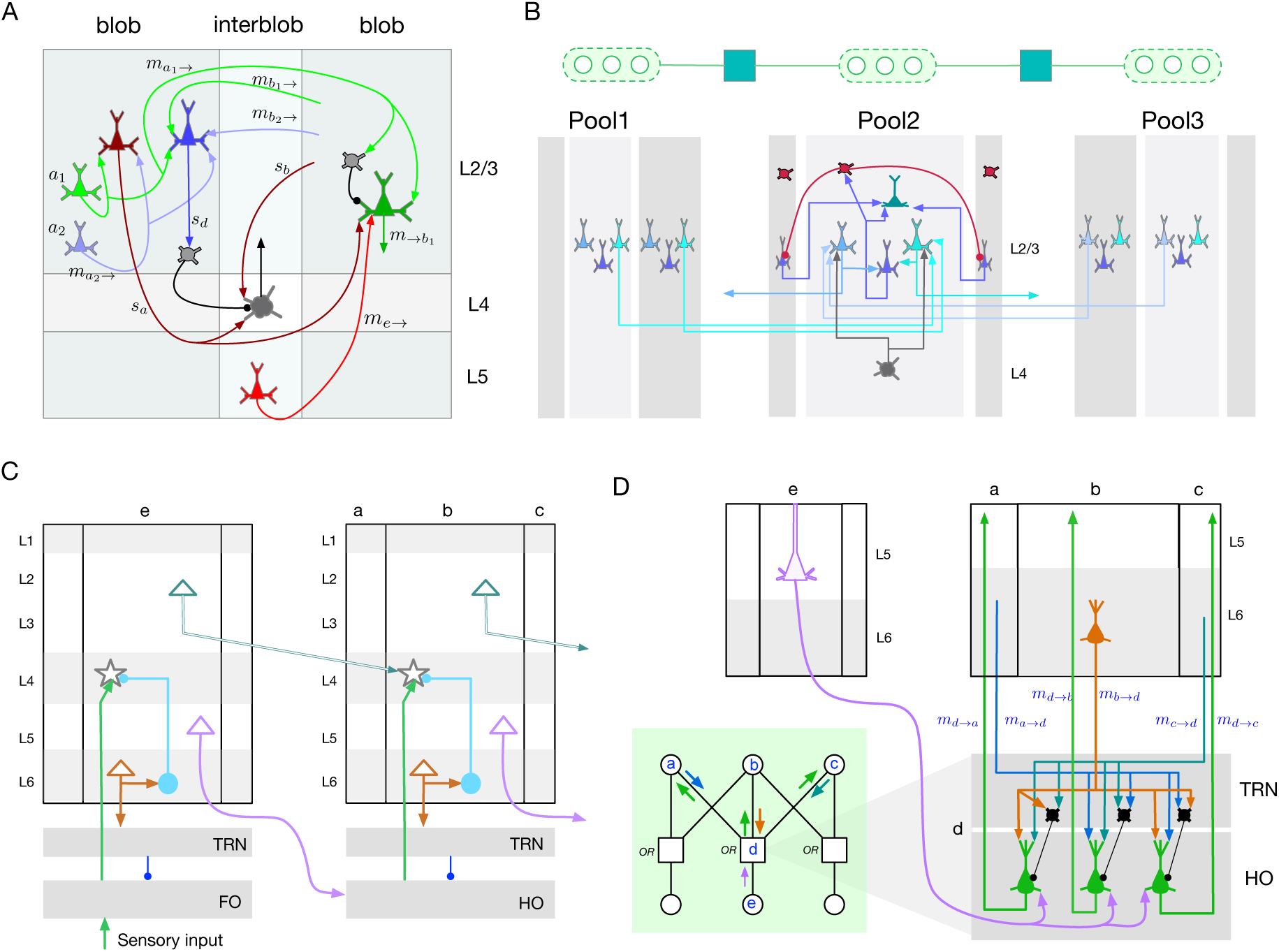
Cortical microcircuit motifs. A. RCN equations predict that interactions between blob and inter-blob columns will include different forms of dendritic integration and gating. B. Detailed microcircuits of contour-contour lateral connections and pooling. The dendrites of the neurons in layer 2/3 encode the compatibility factor (yellow node in the factor graph) between different pools. Columns in a pool also inhibit each other as part of lateral propagation. C. Ineraction between cortical areas V1 and V2, gated through the thalamus. This pattern of connectivity is a repeating motif between different cortical areas. D. (In the green rectangle: factor graph showing the interaction between corticla columns *a, b, c* in V2, with cortical column *e* in V1, through the factor *d*.) Thalamic relay and TRN microcirucit predicted by explaining away computations in RCN, and its connections to child and parent cortical columns. A feed-forward pathway originating in V1 Layer 5 projects to the relay cells in HO thalamus, which are gated by inhibitory TRN cells based on excitatory feedback projections from layer 6 in V2.

### 3.3. Layer 2/3: Pooling and feed-forward output

A subset of the complex pyramidal cells in layer 2/3 [46,47] match the pooling operation in RCN. Pooling, a special case of which is pooling over translations of the same feature, gives invariance to local transformations. In general, pools are flexible and specified non-parameterically through the specific features they connect to. In the feed-forward pass, the POOL factor nodes (Fig 4A, annotation 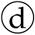) implement a maximum operation over the input messages, and produce the `FF_pool` output (Fig 5 C).

The green neurons annotated 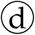) in layer 2/3 or Fig 6 B compute the pooling output in Bio-RCN. Multiple clones of this neuron, all with the same bottom-up inputs, participate in different higher-level features. The pool neurons in a particular column represent the pools that are ‘centered’ at that column. Pooling neurons need to receive trans-columnar inputs from features that are transversal, for example, different translation of a horizontal line. Similar to contour continuity laterals, connections for pooling need to integrate information from multiple columns. The pyramidal neurons in layer 2/3 that receive inputs laterally from other feature columns are ideally suited for performing these computations. The outputs of pooling neurons `FF_pool` is the direct feed-forward output of the region that is sent to the next level, and will be considered in detail in a subsequent section.

### 3.4. Layer 1: Feedback connections from higher-levels

Feedback messages convey top-down certainty about the pools in lower-levels using a scalar corresponding to each pool (`FB_pool` messages in Fig 4A). In Bio-RCN (Fig 6), these feedback lines rise to layer 1 and extend horizontally [13,48,49]. Pyramidal neurons in different columns and laminae can access these messages by extending their distal dendrites vertically into layer 1.

### 3.5. Layer 2/3 or Layer 5: Feedback based unpooling

Unpooling is the reverse operation of pooling, where the evidence of a parent feature being ON is distributed to all its component pools at lower level (Fig 4 A, annotation 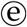). Neurons in a cortical implementation (annotated 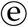 in layer 5 in Fig 6 B) will need to receive feedback information through their apical dendrites, send their outputs across columns to all the features that are part of the pool, and terminate in laminae that are involved in the next stage of feedback processing. A class of neurons in layer 3 and layer 5 both match the requirements for this. The descending projections from layer 2/3 to layer 5 is a classical pathway that has been confirmed in multiple studies [50,51], and is known to control the gain of layer-5 outputs [52,53]. Alternatively, the same computations could be implemented in a class of layer 5 pyramidal neurons with similar connectivity constraints [32].

### 3.6. Layer 5: Lateral propagation with top-down inputs

Bio-RCN predicts that a set of neurons in layer 5 implements lateral propagation similar to the layer 2/3 pyramidal neurons. Feedback computations of contour-continuity laterals are similar to that of feed-forward lateral computations, except for the addition of top-down information. Top-down messages act as a priors on the pools at the lower level, and determine which pools in the children are ON/OFF. The specific feature column that is to be turned ON within a pool is then determined as the one most compatible with its neighboring pools, based on lateral message passing (annotation 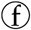 in Fig 4 A and Fig 6 B).

Layer 5 pyramidal cells match the constraints for this because they have long-range lateral arborizations within layer 5 [49,50], and can receive feedback information through their apical dendrites in layer 1 or layer 3 [32,54]. Layer 2/3 pyramidal neurons are a direct source of excitatory input to layer 5, and the cortico-cortical pyramidal neurons in layer 5 A/B are know to have lateral arborizations similar to that of layer 2/3 pyramidal cells, matching the requirements for these being a copy of the contour-continuity lateral connections [14]. Although data detailed data about layer-5 inhibition fields are missing, our mapping predicts that these layer 5 neurons will also have inhibition fields as those suggested for layer 2/3 neurons [45].

### 3.7. Layer 5: Pool Belief as a feed-forward output through the thalamus

RCN provides an explanation for the logic of parallel feed-forward pathway in addition to the direct cortico-cortical feedforward pathway originating from layer 2/3 [55,56]: the cortico-thalamo-cortical pathway provides a mechanisms for explaining away, where thalamus implements the gray OR factor annotated 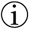 in 4A (equations in Fig 5E). Explaining away is a form of gating mechanism where feed-forward messages are affected by feedback messages. The same neuronal apparatus that supports explaining away can also be used for other forms of gating that includes top-down object based attention (hard explaining away), suppression, or graded gating. To be consistent with a direct feed-foward pathway that conveys evidence about pools, the thalamic feed-forward pathway needs to be about pools as well. The layer-5 pyramidal neurons projecting to the thalamus also receive feedback information from layer 1. Therefore, the feed-forward message through the thalamus is likely to be the belief in pools, which combines both feed-forward and feedback evidence.

The purple outlined neuron in Fig 6 B shows the proposed cortical implementation. The pool likelihood neurons that project from layer 2/3 for a direct feed-forward connection (green neurons marked 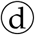 in Fig 6B), also synapses in layer 5. A subset of pyramidal neurons in layer 5 that receive feedback information through their dendrites in layer 1 can compute pool beliefs by combining input from this pool likelihood axon branch. According to Bio-RCN mapping, these layer 5 neurons project to the thalamus, where they take part in explaining away computations and gating, before projecting to the next level in the hierarchy. Belief propagation allows for sending either likelihoods or beliefs as feed-forward messages [57] because the next level in the hierarchy can derive likelihoods from the belief message by subtracting out the feedback message originating from that region. In Bio-RCN these computations are implemented as part of the explaining away in the thalamus, and will be considered in more detail in a subsequent section.

In the current version of RCN, a child region cannot turn on pools that are not turned on by the parent region, a constraint that we expect to be relaxed in future versions to support more dynamic assemblies based on lateral evidence. This would require projections from the lateral computations, shown as dotted axons from the blue pyramidal neurons in layer 5 of Fig 6. The need to turn ON pools not supported by top-down lends further support to need for the existence of a pool belief neuron.

### 3.8. Layer 5: Calculation of feature beliefs

Feature belief is the summary output of a column on its status (ON or OFF) incorporating all available evidence (`BEL_feat` in 4(C)). The final step in this computation is a summation of evidence from feature-copies that participate in different lateral connections. A subset of neurons in layer 5, which are downstream from the lateral computation circuit (green neurons annotated 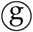 in layer 5 of Fig 6) match the connectivity requirements for this computation [52]. These neurons project to subcortical circuits [58], arguably because it is advantageous to drive actions from beliefs.

### 3.9. Layer 6: Feedback message to child regions

Connections of layer 6 neurons [51] are consistent with the anatomical requirement for computations of RCN feedback messages from one level to its children (annotation h in Fig 4 A). Feedback messages for child regions are the reverse of the feed-forward computation where the belief of a feature is unpacked into its components and sent to the corresponding child regions. The three brown neurons annotated 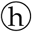 in layer 6 of Fig 6 correspond to the feedback messages to the three components of the feature represented by this column. Many feed-forward thalamo-cortical axons that synapse in layer 4, also send axon branches to layer 6 [49]. The details of this layer 6 projection are unknown, but RCN can suggests one explanation where these projections help to subtract out the feed-forward influence from the feedback projections.

### 3.10. Functional role of thalamus: Explaining away, gating, top-down attention, and binding

The functional logic of the thalamic pathway has been a source of enduring mystery in neuroscience [59]. Anatomical data show two feed-forward circuits: a direct cortico-cortical connection from layer 2/3, and an indirect cortico-thalamo-cortical connection from layer 5 [55,60,61]. The thalamus also receives feedback connections from layer-6 neurons in the higher level [55,62,63]. The higher-level neurons that send feedback projections from layer-6 also project back to layer-4 of the same region via an inhibitory circuit as shown in Fig 7C. Feedback projections from layer 6, project to the TRN of the thalamus and makes precise connections that modulate the feed-forward pathway.

RCN message passing offers an integrated explanation for functional role of the thalamus, including many of the details of the thalamic circuits, and connections spanning multiple cortical regions. The gray-colored OR factors in RCN (Fig 2 (C)) encode how multiple features interact while generating the same lower-level components. Such multi-cause interactions produce explaining away effects during inference where the strength of one hypothesis is diminished when the same cause can be explained by the presence of another. These OR factors are also the interfaces between sensory input and the first level, and between different levels. According to RCN, the thalamus implements this OR factor and the associated explaining away mechanisms and sits at the interface between two levels (Fig 4(A)).

Known details of the precise connections to thalamus and TRN are consistent with this proposal [63]. Consider the PGM fragment in Fig 7 D, where the nodes *a, b, c* correspond to features at a higher level (V2) and nodes *e, f, g* correspond to pools at a lower level (V1). The factor-nodes (e.g., node *d*) in this computation implement explaining away, where the feed-forward messages are affected by feedback messages according to equations in Fig 5E. Explaining away requires feed-forward inhibition [64], and our mapping suggests that thalamic reticular nucleus (TRN) implements this function as shown in Fig 6B and Fig 7D. Feedback connections originating from layer 6 project to the inhibitory cells in TRN to modulate the axons of the driving feed-forward connections originating from sensory input or layer-5 of the region below, and passing through TRN. These feedback connections are reciprocal in the sense that they respect the topography of the feed-forward connections [63], and they cross connect with feed-forward axons[65], matching the requirement for explaining away [64]. TRN is organized retinotopically enabling it to receive spatially localized feedback information and to influence thalamic relay cells in spatially specific ways [66]. A detailed view of this circuit between two cortical regions is shown in Fig 7D. According to Bio-RCN, the explaining away circuit shown will be a motif that repeats in LGN, pulvinar, and in general when two cortical areas interact through the thalamus.

Attention gating of various kinds are special cases of explaining away computation, and the need to support these additional mechanisms provide a rationale for the explaining away factor being implemented in a separate structure that can act as switchboard for different control signals [66]. One example of a special case is top-down attention on an object or on a set of features. In the factor graph, this corresponds to setting the parent nodes to be ON, propagating that information everywhere, and then letting the feed-forward flow of information being affected by that. In the neural implementation this will require sustaining the activity of neurons turned on by the top-down attention, which is something that thalamus could support. Top-down attention to a set of features, or colors, can operate in a similar manner. Lesions in the pulvinar, a higher-order thalamic nucleus, are implicated in deficits in binding object color to object shape using top-down attention [66]. Other special cases include turning OFF the attention to a particular object of set of features, or reasoning about occlusion and amodal completion.

### 3.11. Parallel feed-forward pathways

RCN offers rationale for two feed-forward pathways between cortical regions as observed in biology [63]. The direct cortico-cortical pathway originates in layer 2/3 and projects to layer 4 of the next level, and the indirect cortico-thalo-cortical path originates from layer 5 and is gated through the thalamus and includes explaining away (Fig 7C). One special case of explaining away computations is when all parent nodes are assumed to be OFF. In this case, the bottom-up messages are copied to all parent nodes without any feed-forward inhibition. In practically used implementations of RCN, the first forward pass of evidence operates in this manner as well, and serves to obtain an approximate but fast inference over higher-level features. For an animal, having a fast, prior-free, feed-forward pathway could be advantageous because it can alert the animal to novel, out-of-context situations. The pathway that goes through the thalamus includes explaining away and attention control, and could be slower in switching to a rapid input-driven change. The inhibitory projection from layer 6 to layer 4 might be an approximate version of this explaining away circuit as well, providing a faster, but approximate, explaining away.

A second justification, consistent with the first, can be offered by the need for keeping imaginations separate from reality. The perceptual apparatus of RCN can be used by a cognitive system [67] for visual imagery as part of planning, exploration, and conceptual thinking. A direct feed-forward pathway could offer a mechanism for quickly snapping out of imagination to synchronize back with reality. Mixing imagery with evidence could be behind the hallucinations in schizophrenia, a topic previously explored using probabilistic models [68–70]. The detailed visual circuit offered by RCN could bring further precision to such investigations.

### 3.12. Interaction between shape and appearance (blobs and inter-blobs)

The factorized contour-surface representation of RCN, which enables generalization to novel combinations of shape and appearance, offers an explanation for the existence of blobs and inter-blobs [33,71] in the primary visual cortex, and predicts circuit-level details of their interactions. The contour-surface factor in RCN (Fig 4(C)) encodes a three way interaction: it enforces continuity of surface properties (eg., color, texture, surface angle, etc.) between adjacent surface patches except when interrupted by an intervening contour (Fig 5 D). Message passing through this three-way factor enhances the support for contours when the surface patches are discontinuous and vice versa. RCN uses an efficient reparametrization of these message computations that avoids the quadratic set of interconnections required in a naive implementation (See supplementary material for details). Fig 6B shows the positioning of the blob inter-blob interactions within the overall cortical column. Detailed predicted circuit of the interaction between interblob columns and blobs are shown in Fig 7 A, and it utilizes multiple excitatory neuron types, inhibition, and dendrite-specific gating mechanisms to achieve the required computation.

## 4. Explanations for visual phenomena

RCN was successful in reproducing and explaining three different visual phenomena – subjective contours, neon color spreading, and occlusion versus deletion effect – to the details of neuron-level activations and their dynamics in different cortical laminae and columns. Notably, all these phenomena are explained as the byproduct of inference in a model that was constructed and learned for parsing visual scenes.

### Subjective contours

RCN that is trained to recognize regular shapes ‘hallucinates’ illusory contours in visual stimuli (Fig 8 A & B) that produce subjective contour perception in people. Subjective contours is an illusion because people see contours that are not present in the input. RCN produce these hallucinations as a natural byproduct of ‘inference to best explanation’ — according to the model, evidence in the whole of the image is best explained by hallucinating these contours. In Fig. 8 A & B, the contours shown in magenta and green constitute the MAP solution at the lowest level of the network, where magenta color indicates contours that exist according to RCN beliefs although they are not supported by local bottom-up evidence.

**Figure 8:**
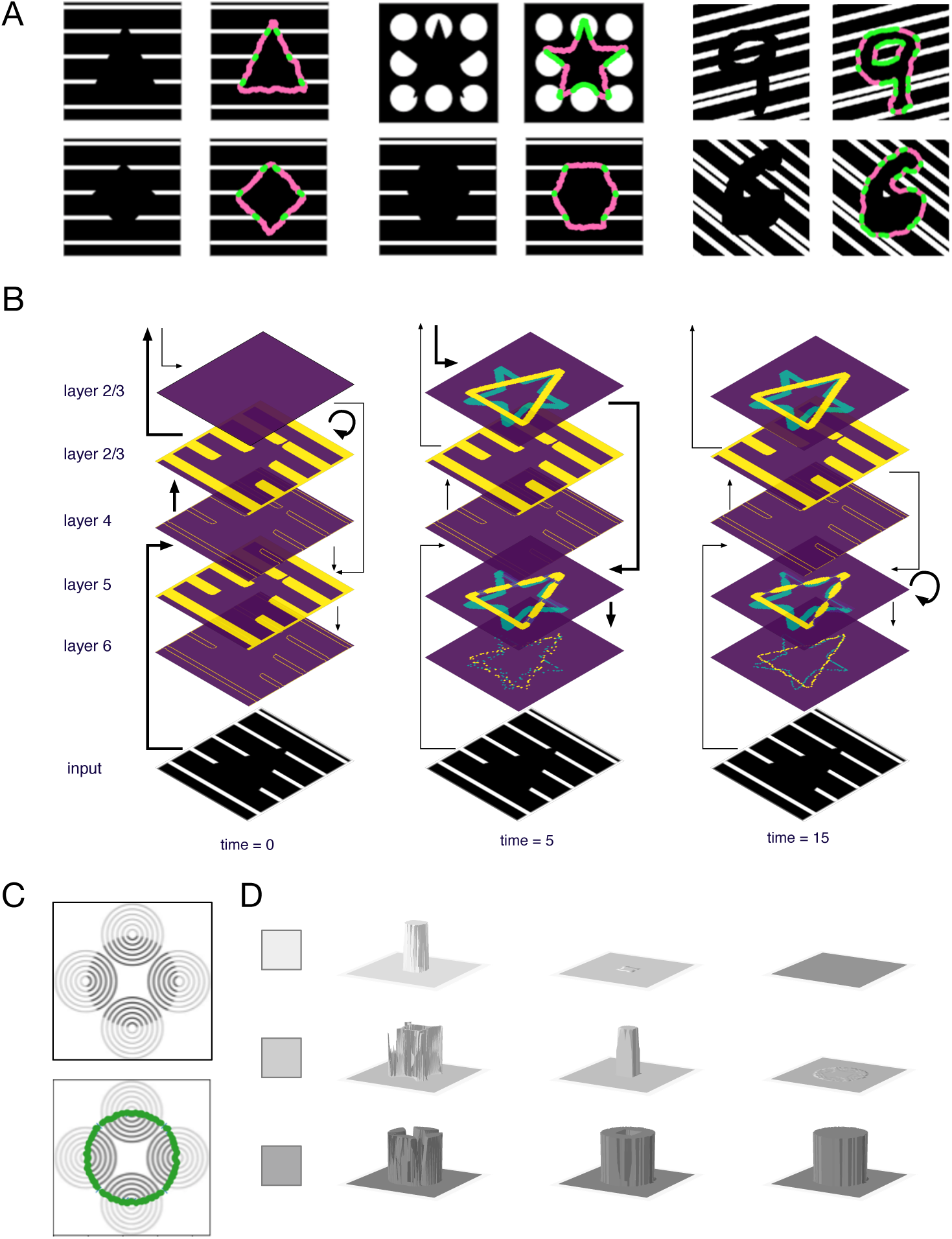
Subjective contours and neon-color spreading. A) RCN MAP inference results for stimuli that induce subjective contours in humans. Green: Top-down imaginations with bottom-up evidential support. Magenta: Contour hallucinations. B) Layer-wise dynamics of subjective contour formation in Bio-RCN. C) Stimulus for neon color spreading, and the subjective contour backtrace. D) Dynamics of color spreading inside the surface, for three different colors.

The dynamics of subjective contour effects in RCN produce a qualitative match to those observed in biology (Fig 8(C)). Physiological results report evidence for neurons in V1 responding to the illusory contour, albeit with a delay compared to the neurons responding to real contours [16,72,73]. The temporal dynamics of neuronal responses to subjective contours can be readily understood from the schedule of message propagation. During the forward pass, the features have only local evidence, and hence the neurons in blank spaces do not respond. Once forward pass identifies a potential global percept, that information flows down in the top-down messages to affect the beliefs in lower level nodes to turn ON some features that were previously OFF.

Bio-RCN’s delineation of feed-forward, lateral, and feedback propagation in shape perception to laminar and columnar precision can help clarify long-standing debates on whether shape perception reduces or enhances activity in the primary visual cortex [16,72,74–76]. The observation that neural activity is increased in regions with no bottom-up evidence [76], is explained by top-down and lateral evidence that is consistent with the global percept, and Bio-RCN’s prediction on this are precise to the level of the specific neuronal populations in laminae and columns. Concurrently, the observation that activity in regions that receive bottom up input consistent with top-down prediction [76] is also explained by Bio-RCN. In the forward pass, all the potential pool and lateral participation of the presence of a feature are active, and feedback and lateral propagation will reduce these activities to those are consistent with the global percept. Note that this explanation is different from that of ‘predictive coding’ models that require predictions to be subtracted from the input. Bio-RCN also makes lamina-specific predictions that top-down attention will reduce the activity in background elements that were active in the forward pass, which might be useful for designing further experiments on the differentiated roles of different laminae in shape perception [77].

### Neon color spreading

Inference in RCN reproduced the effect of neon-color spreading, the perception of an illusory surface with an illusory color [78], and its cortical implementation predicts circuit dynamics at the neuronal level (Fig. 8 D). The suggested mechanism behind these effects is the interplay between boundary completion and surface filling-in in visual cortex [17]. Notably, the filling in of the illusory surface respects the boundaries of the illusory contours. The neon color spreading effect is a natural byproduct of the dynamics of MAP inference in RCN. Three-way interactions in the contour-surface CRF encourages continuity between adjacent surface patches unless the intervening contour node is turned ON. A forward pass through this model produces approximate edge and surface responses, and leads to the selection of ‘circle’ as the best top-level hypothesis. The backward pass, which is based on selecting the most active hypothesis at the top level of the contour hierarchy, will then enforce the corresponding contour discontinuities on the surface CRF. Similar to the subjective contour effects described earlier, the stimulus shown in Fig 8 D has sufficient local edge evidence to support a circle as the top level hypothesis in the RCN contour hierarchy, and to hallucinate the missing contours of the circle. The top-down partial MAP configuration for contours of the circle, including the hallucinated portions, then influences the propagation in the CRF. The discontinuity imposed by the top-down contours will propagates in the CRF with further message passing to create the fill-in effect with the dynamics shown in Fig 8 D.

The contour formation (Fig 8B) and the surface filling dynamics in (Fig 8D) Bio-RCN match the dynamics and laminar profiles of figure-ground segregation observed in cortical layers [79]. Matching biological observations, layer-4 neurons are activated first, followed by layer 2/3 in the feed-forward pass. Labeling of the perceived figure with enhanced activity occurs in layers 2/3 and layer 5, and region-filling is determined by the layers that receive feedback information from higher level [77,79,80]. Taken together, neon-color spreading in Bio-RCN support the observation that object completions involve feedback loops that integrates contour interpolation with surface filling in [81].

### Occlusion versus deletion

RCN also reproduces the psychophysics observation [82] that humans are much better at detecting objects under occlusion than the same objects where the same regions have been deleted instead of occluded, while keeping identical visible portions (Fig. 9. Occlusion reasoning performance of RCN was already demonstrated in [10], and the RCN generative model offers the reasoning for why partially occluded images are easier to recognize compared to their partially deleted counterparts. Deletion of the parts of an object is absence of evidence for those parts. When those same parts are missing due to occlusion, the model can explain away the absence of evidence as occlusion. Mechanistically, the portions that are deleted will contribute negative evidence to the overall hypothesis if there is no occlusion to explain their absence. Explaining away during occlusion reasoning will convert those negative evidences to ‘uncertain evidence’ (log-likelihood = 0). The dynamics of occlusion reasoning and filling in have limited studies in neuroscience [83–85], and predictions from Bio-RCN might be a useful tool in further explorations.

**Figure 9:**
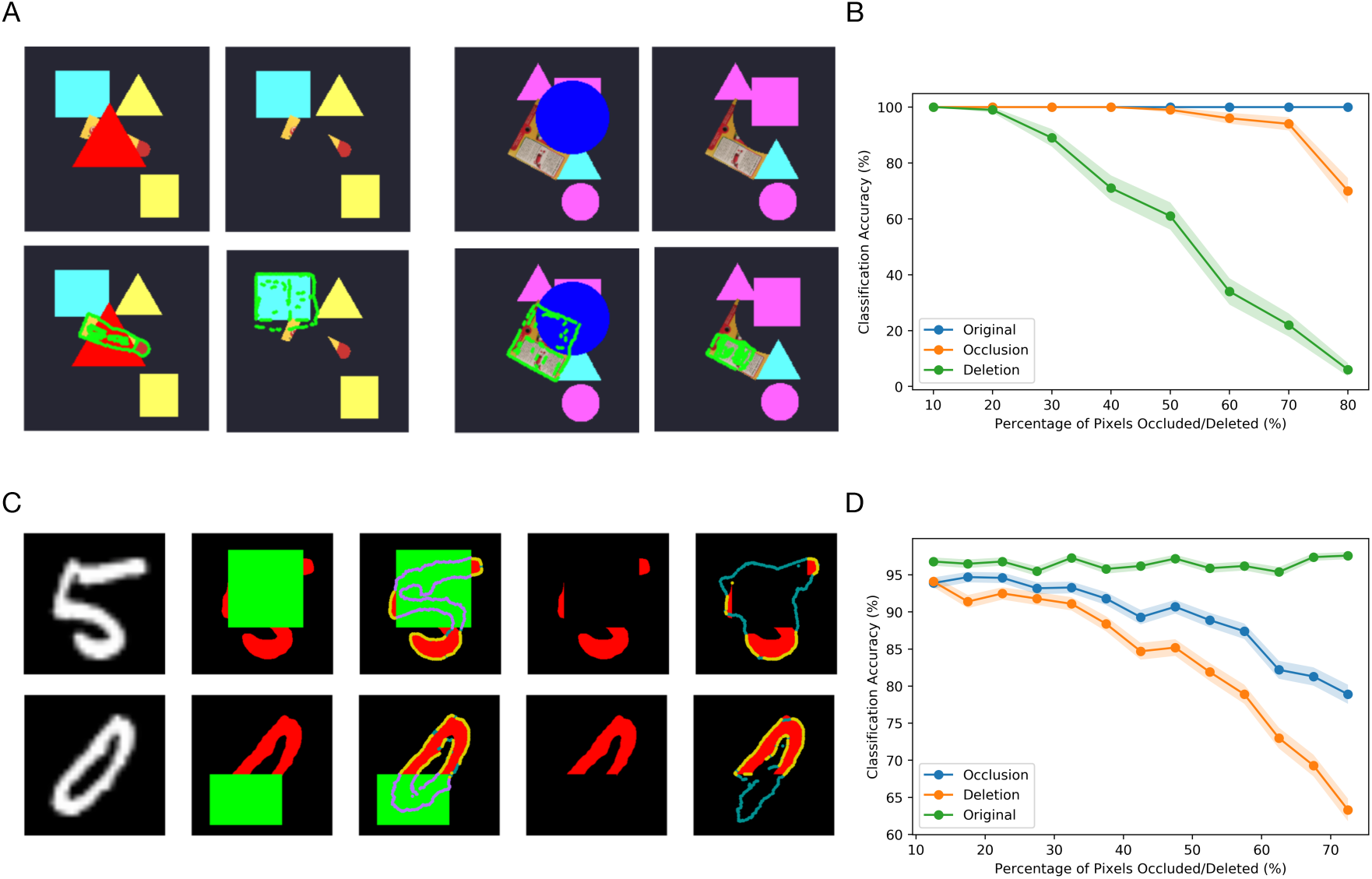
Occlusion vs deletion. A) Humans have a harder time perceiving objects with deleted portions compared to the case where the same portions are occluded. Clockwise in each each set of 4 images: (1) Object with occlusion, (2) the same object with occluded portion deleted, (3) RCN MAP inference for occlusion case, (4), RCN MAP inerence for deletion. B) Recognition accuracy for occlusion vs deletion as a function of the amount of missing evidence C) Occlusion vs deletion for MNIST digits. In each row left to right, original digit, occluded digit, MAP inference overlaid on occlusion, digit with occluded portions deleted, MAP inference with deletion. D) Same as B, for MNIST digits.

## 5. Discussion

Here, we derived a detailed and functional mathematical theory of cortical and thalamic microcircuits underlying visual perception. The approach we followed can be construed as a general methodology by which we can seek to understand cortical function. The first step in this methodology is to triangulate between cortical data, properties of the world, and computational principles to build algorithmic models that are tested on real-world tasks. The second step considers the algorithmic model in conjunction with anatomical and physiological data to derive detailed and functional microcircuit-level models. The advantage of this approach is that the derived circuit is methodology-constrained to a coherent functional model that cuts across different organizational levels – cortical regions, thalamus, cortical columns, cortical laminae, neuron types, and dendrites.

Our approach is in contrast to large scale systems neuroscience efforts gathering large amounts of data on cortical connectivity [5,86,87]. While more data is definitely useful, purely data-driven simulations is an unlikely path to understanding cortical function. We believe that the neuroscience-guided theoretical approach we use here is a necessary complement to data-gathering and experimental approaches.

Message-passing inference in RCN, the algorithmic base for the microcircuit model, is more general and more effective than typical predictive coding [11] accounts. Complex problems like perception might require more general non-parameteric inference compared to predictive coding which makes restrictive linear/Gaussian assumptions. Our experiments demonstrate that various accounts of response reduction and enhancement can be accounted using our more general generative model without the restrictive assumptions of predictive coding [88].

While known neuroanatomy and physiology impose constraints on cortical implementation of a set of inference equations, several variations of the same implementation are possible because neurons can be moved to different laminae, or a computation can be accomplished in dendrites instead of neurons. We consider a few such variations in the supplementary material. Furture research could explore this hodology in more detail, and combine it with wiring and metabolic constraints. The model made several predictions about potential microcircuit connections. Further neuroscience experiments could test the veracity of these predictions and suggest alternative implementations. Predictions from our circuit model could also help further investigations of neurological and psychiatric conditions [68,70].

RCN currently is a model of the ventral visual pathway. While many aspects of the derived microcircuit are likely to generalize to other visual pathways, some aspects like blob-interblob interactions are likely to be specific to the ventral pathway. RCN could potentially be extended to utilize temporal information for pooling [89], and to utilize binocular disparity for depth perception. In future work, we intend to explore those extensions and study how they might result in a more comprehensive model of visual cortical microcircuits.

We started with an algorithmic AI model that was inspired by observations from neuroscience, and showed how that model can be used for deriving neuroscience insights. Such interactions between artificial intelligence and neuroscience research are likely to simultaneously help accelerate progress in both fields towards two ultimate goals: building artificial general intelligence, and understanding the brain.

## Acknowledgments

We thank Adam Marblestone, Alex Naka, S.P. Arun, Doris Tsao, and Drew Linsley for their insightful and patient comments after reviewing early versions of this manuscript. Those helped to greatly improve the manuscript.

## References

1. Adesnik, H.; Naka, A. Cracking the Function of Layers in the Sensory Cortex, 2018. doi: 10.1016/j.neuron.2018.10.032.

2. Emiliani, V.; Cohen, A.E.; Deisseroth, K.; Häusser, M. All-optical interrogation of neural circuits, 2015. doi: 10.1523/JNEUROSCI.2916-15.2015.

3. Luo, L.; Callaway, E.M.; Svoboda, K. Genetic Dissection of Neural Circuits: A Decade of Progress, 2018. doi: 10.1016/j.neuron.2018.03.040.

4. Marr, D. Vision: A Computational Investigation into the Human Representation and Processing of Visual Information.; MIT Press, 1982.

5. Markram, H.; Muller, E.; Ramaswamy, S.; Reimann, M.W.; Abdellah, M.; Sanchez, C.A.; Ailamaki, A.; Alonso-Nanclares, L.; Antille, N.; Arsever, S.; Kahou, G.A.A.; Berger, T.K.; Bilgili, A.; Buncic, N.; Chalimourda, A.; Chindemi, G.; Courcol, J.D.; Delalondre, F.; Delattre, V.; Druckmann, S.; Dumusc, R.; Dynes, J.; Eilemann, S.; Gal, E.; Gevaert, M.E.; Ghobril, J.P.; Gidon, A.; Graham, J.W.; Gupta, A.; Haenel, V.; Hay, E.; Heinis, T.; Hernando, J.B.; Hines, M.; Kanari, L.; Keller, D.; Kenyon, J.; Khazen, G.; Kim, Y.; King, J.G.; Kisvarday, Z.; Kumbhar, P.; Lasserre, S.; Le Bé, J.V.; Magalhães, B.R.; Merchán-Pérez, A.; Meystre, J.; Morrice, B.R.; Muller, J.; Muñoz-Céspedes, A.; Muralidhar, S.; Muthurasa, K.; Nachbaur, D.; Newton, T.H.; Nolte, M.; Ovcharenko, A.; Palacios, J.; Pastor, L.; Perin, R.; Ranjan, R.; Riachi, I.; Rodríguez, J.R.; Riquelme, J.L.; Rössert, C.; Sfyrakis, K.; Shi, Y.; Shillcock, J.C.; Silberberg, G.; Silva, R.; Tauheed, F.; Telefont, M.; Toledo-Rodriguez, M.; Tränkler, T.; Van Geit, W.; Díaz, J.V.; Walker, R.; Wang, Y.; Zaninetta, S.M.; Defelipe, J.; Hill, S.L.; Segev, I.; Schürmann, F. Reconstruction and Simulation of Neocortical Microcircuitry. Cell 2015, 163, 456–492. doi: 10.1016/j.cell.2015.09.029.

6. Lee, T.; Mumford, D. Hierarchical Bayesian inference in the visual cortex. JOSA A 2003, 20, 1434–1448.

7. Friston, K. The free-energy principle: a unified brain theory? Nature reviews. Neuroscience 2010, 11, 127–138. doi: 10.1038/nrn2787.

8. Jones, M.; Love, B.C. Bayesian Fundamentalism or Enlightenment? On the explanatory status and theoretical contributions of Bayesian models of cognition. Behavioral and Brain Sciences 2011, 34, 169–188. doi: 10.1017/S0140525X10003134.

9. Grossberg, S.; Pearson, L.R. Laminar cortical dynamics of cognitive and motor working memory, sequence learning and performance: toward a unified theory of how the cerebral cortex works. Psychological review 2008, 115, 677–732. doi: 10.1037/a0012618.

10. George, D.; Lehrach, W.; Kansky, K.; Lázaro-Gredilla, M.; Laan, C.; Marthi, B.; Lou, X.; Meng, Z.; Liu, Y.; Wang, H.; Lavin, A.; Phoenix, D.S. A generative vision model that trains with high data efficiency and breaks text-based CAPTCHAs. Science 2017. doi: 10.1126/science.aag2612.

11. Bastos, A.M.; Usrey, W.M.; Adams, R.A.; Mangun, G.R.; Fries, P.; Friston, K.J. Perspective Canonical Microcircuits for Predictive Coding. Neuron 2012, 76, 695–711. doi: 10.1016/j.neuron.2012.10.038.

12. Oberlaender, M.; De Kock, C.P.; Bruno, R.M.; Ramirez, A.; Meyer, H.S.; Dercksen, V.J.; Helmstaedter, M.; Sakmann, B. Cell type-specific three-dimensional structure of thalamocortical circuits in a column of rat vibrissal cortex. Cerebral Cortex 2012, 22, 2375–2391. doi: 10.1093/cercor/bhr317.

13. Harris, K.D.; Mrsic-Flogel, T.D. Cortical connectivity and sensory coding, 2013. doi: 10.1038/nature12654.

14. Harris, K.D.; Shepherd, G.M. The neocortical circuit: Themes and variations, 2015. doi: 10.1038/nn.3917.

15. Lee, T.S.; Nguyen, M. Dynamics of subjective contour formation in the early visual cortex. Proc Natl Acad Sci U S A 2001, 98, 1907–1911. doi: 10.1073/pnas.031579998.

16. Meng, M.; Remus, D.A.; Tong, F. Filling-in of visual phantoms in the human brain. Nature Neuroscience 2005, 8, 1248–1254. doi: 10.1038/nn1518.

17. Grossberg, S. Filling-In the Forms: Surface and Boundary Interactions in Visual Cortex. In Filling-In: From Perceptual Completion to Cortical Reorganization; Oxford University Press, 2009. doi: 10.1093/acprof:oso/9780195140132.003.0002.

18. Pearl, J. A Constraint Propagation Approach to Probabilistic Reasoning. In Uncertainty in Artificial Intelligence 2; Kanal, L.F.; Lemmer, J.F., Eds.; North-Holland, 1988.

19. Frey, B.J.; MacKay, D.J.C. A Revolution: Belief Propagation in Graphs with Cycles. Advances in Neural Information Processing Systems; Jordan, M.I.; Kearns, M.J.; Solla, S.A., Eds. The MIT Press, 1998, Vol. 10.

20. Yuille, A.; Kersten, D. Vision as Bayesian inference: analysis by synthesis? Trends in Cognitive Sciences 2006, 10, 301–308. doi: 10.1016/j.tics.2006.05.002.

21. Poirazi, P.; Papoutsi, A. Illuminating dendritic function with computational models. Nature Reviews Neuroscience 2020, pp. 1–19. doi: 10.1038/s41583-020-0301-7.

22. Mountcastle, V.B. Perceptual Neuroscience: The Cerebral Cortex; Harvard University Press: Cambridge, MA, 1998.

23. Li, Y.; Lu, H.; Cheng, P.L.; Ge, S.; Xu, H.; Shi, S.H.; Dan, Y. Clonally related visual cortical neurons show similar stimulus feature selectivity. Nature 2012, 486, 118–121. doi: 10.1038/nature11110.

24. Ohtsuki, G.; Nishiyama, M.; Yoshida, T.; Murakami, T.; Histed, M.; Lois, C.; Ohki, K. Similarity of visual selectivity among clonally related neurons in visual cortex. Neuron 2012, 75, 65–72. doi: 10.1016/j.neuron.2012.05.023.

25. Cadwell, C.R.; Scala, F.; Fahey, P.G.; Kobak, D.; Mulherkar, S.; Sinz, F.H.; Papadopoulos, S.; Tan, Z.H.; Johnsson, P.; Hartmanis, L.; Li, S.; Cotton, R.J.; Tolias, K.F.; Sandberg, R.; Berens, P.; Jiang, X.; Tolias, A.S. Cell type composition and circuit organization of clonally related excitatory neurons in the juvenile mouse neocortex. eLife 2020, 9. doi: 10.7554/elife.52951.

26. Dedieu, A.; Gothoskar, N.; Swingle, S.; Lehrach, W.; Lázaro-Gredilla, M.; George, D. Learning higher-order sequential structure with cloned HMMs. arXiv preprint 1905.00507 2019.

27. Rikhye, R.V.; Gothoskar, N.; Guntupalli, J.S.; Dedieu, A.; Lázaro-Gredilla, M.; George, D. Learning cognitive maps as structured graphs for vicarious evaluation. bioRxiv 2020, [https://www.biorxiv.org/content/early/2020/06/24/864421.full.pdf]. doi: 10.1101/864421.

28. Medini, P. Experience-dependent plasticity of visual cortical microcircuits, 2014. doi: 10.1016/j.neuroscience.2014.08.022.

29. Cadwell, C.R.; Scala, F.; Fahey, P.G.; Kobak, D.; Sinz, F.H.; Johnsson, P.; Li, S.; Cotton, R.J.; Sandberg, R.; Berens, P.; Jiang, X.; Tolias, A.S. Cell type composition and circuit organization of neocortical radial clones. bioRxiv 2019, p. 526681. doi: 10.1101/526681.

30. Shipp, S. Visual Processing: The odd couple. Current Biology 1995, 5, 116–119. doi: 10.1016/S0960-9822(95)00029-7.

31. Lund, J.S.; Angelucci, A.; Bressloff, P.C. Anatomical substrates for functional columns in macaque monkey primary visual cortex. Cerebral cortex (New York, N.Y. : 1991) 2003, 13, 15–24.

32. Thomson, A.M.; Lamy, C. Functional maps of neocortical local circuitry. Frontiers in neuroscience 2007, 1, 19–42. doi: 10.3389/neuro.01.1.1.002.2007.

33. Sincich, L.C.; Horton, J.C. THE CIRCUITRY OF V1 AND V2: Integration of Color, Form, and Motion. Annual Review of Neuroscience 2005, 28, 303–326. doi: 10.1146/annurev.neuro.28.061604.135731.

34. Callaway, E. Local circuits in primary visual cortex of the macaque monkey. Annual review of neuroscience 1998, 21, 47–74.

35. Ben-Shahar, O.; Zucker, S. Geometrical Computations Explain Projection Patterns of Long-Range Horizontal Connections in Visual Cortex. Neural Computation 2004, 16, 445–476. doi: 10.1162/089976604772744866.

36. Zucker, S.W.; Wagemans, J. Border Inference and Border Ownership Border Inference and Border Ownership: The Challenge of Integrating Geometry and Topology. Oxford Handbook of Perceptual Organization 2014. doi: 10.1093/oxfordhb/9780199686858.013.020.

37. Samonds, J.M.; Zhou, Z.; Bernard, M.R.; Bonds, A.B. Synchronous activity in cat visual cortex encodes collinear and cocircular contours. Journal of neurophysiology 2006, 95, 2602–16. doi: 10.1152/jn.01070.2005.

38. Karube, F.; Kisvárday, Z.F. Axon topography of layer IV spiny cells to orientation map in the cat primary visual cortex (Area 18). Cerebral Cortex 2011, 21, 1443–1458. doi: 10.1093/cercor/bhq232.

39. Binzegger, T.; Douglas, R.J.; Martin, K.A. A quantitative map of the circuit of cat primary visual cortex. Journal of Neuroscience 2004, 24, 8441–8453. doi: 10.1523/JNEUROSCI.1400-04.2004.

40. Angelucci, A.; Levitt, J. Circuits for local and global signal integration in primary visual cortex. The Journal of … 2002.

41. Cossell, L.; Iacaruso, M.F.; Muir, D.R.; Houlton, R.; Sader, E.N.; Ko, H.; Hofer, S.B.; Mrsic-Flogel, T.D. Functional organization of excitatory synaptic strength in primary visual cortex. Nature 2015, 518, 399–403. doi: 10.1038/nature14182.

42. Bannister, P.A. Inter- and intra-laminar connections of pyramidal cells in the neocortex. Neurosci Res 2005, 53, 95–103. doi: 10.1016/j.neures.2005.06.019.

43. Lund, J.; Angelucci, A.; Bressloff, P. Anatomical substrates for functional columns in macaque monkey primary visual cortex. Cerebral Cortex 2003, 13, 15–24.

44. Ko, H.; Cossell, L.; Baragli, C.; Antolik, J.; Clopath, C.; Hofer, S.B.; Mrsic-Flogel, T.D. The emergence of functional microcircuits in visual cortex. Nature 2013, 496, 96–100. doi: 10.1038/nature12015.

45. Adesnik, H.; Bruns, W.; Taniguchi, H.; Huang, Z.J.; Scanziani, M. A neural circuit for spatial summation in visual cortex. Nature 2012, 490, 226–230. doi: 10.1038/nature11526.

46. Hubel, D.H.; Wiesel, T.N. Receptive fields and functional architecture of monkey striate cortex. Journal of Physiology 1968, 195, 215–243.

47. Hirsch, J.A.; Martinez, L.M. Laminar processing in the visual cortical column. Curr Opin Neurobiol 2006, 16, 377–384. doi: 10.1016/j.conb.2006.06.014.

48. Markov, N.T.; Kennedy, H. The importance of being hierarchical, 2013. doi: 10.1016/j.conb.2012.12.008.

49. Thomson, A.M.; Bannister, a.P. Interlaminar connections in the neocortex. Cerebral cortex (New York, N.Y. : 1991) 2003, 13, 5–14.

50. Gilbert, C.D.; Wiesel, T.N. Morphology and intracortical projections of functionally characterised neurones in the cat visual cortex. Nature 1979, 280, 120–125. doi: 10.1038/280120a0.

51. Douglas, R.J.; Martin, K.a.C. Neuronal circuits of the neocortex. Annual review of neuroscience 2004, 27, 419–51. doi: 10.1146/annurev.neuro.27.070203.144152.

52. Quiquempoix, M.; Fayad, S.L.; Boutourlinsky, K.; Leresche, N.; Lambert, R.C.; Bessaih, T. Layer 2/3 Pyramidal Neurons Control the Gain of Cortical Output. Cell Reports 2018, 24, 2799–2807. doi: 10.1016/j.celrep.2018.08.038.

53. Kritzer, M.F.; Goldman-Rakic, P.S. Intrinsic circuit organization of the major layers and sublayers of the dorsolateral prefrontal cortex in the rhesus monkey. Journal of Comparative Neurology 1995, 359, 131–143. doi: 10.1002/cne.903590109.

54. Burkhalter, a. Intrinsic connections of rat primary visual cortex: laminar organization of axonal projections. The Journal of comparative neurology 1989, 279, 171–86. doi: 10.1002/cne.902790202.

55. Sherman, S.M. Thalamus plays a central role in ongoing cortical functioning, 2016. doi: 10.1038/nn.4269.

56. Guillery, R.W.; Sherman, S.M. The thalamus as a monitor of motor outputs. Philos Trans R Soc Lond B Biol Sci 2002, 357, 1809–1821. doi: 10.1098/rstb.2002.1171.

57. Pearl, J. Probabilistic reasoning in intelligent systems: networks of plausible inference; Morgan Kaufman, 1988.

58. Harris, K.D.; Shepherd, G.M. The neocortical circuit: Themes and variations, 2015. doi: 10.1038/nn.3917.

59. Rikhye, R.V.; Wimmer, R.D.; Halassa, M.M. Toward an Integrative Theory of Thalamic Function. Annual Review of Neuroscience 2018, 41, 163–183. doi: 10.1146/annurev-neuro-080317-062144.

60. Sherman, S.M.; Guillery, R.W. The role of the thalamus in the flow of information to the cortex. Philos Trans R Soc Lond B Biol Sci 2002, 357, 1695–1708. doi: 10.1098/rstb.2002.1161.

61. Guillery, R.; Feig, S.; Lozsadi, D. Paying attention to the thalamic reticular nucleus. Trends in Neurosciences 1998, 21, 28–31.

62. Rees, G. Visual Attention: The Thalamus at the Centre? Current Biology 2009/03/10, 19, R213–R214.

63. Sherman, S.M. Functioning of Circuits Connecting Thalamus and Cortex. Comprehensive Physiology 2017, 7, 713–739. doi: 10.1002/cphy.c160032.

64. Lochmann, T.; Ernst, U.A.; Denève, S. Perceptual inference predicts contextual modulations of sensory responses. Journal of Neuroscience 2012, 32, 4179–4195. doi: 10.1523/JNEUROSCI.0817-11.2012.

65. Lam, Y.W.; Sherman, S.M. Functional organization of the somatosensory cortical layer 6 feedback to the thalamus. Cerebral cortex (New York, N.Y. : 1991) 2010, 20, 13–24. doi: 10.1093/cercor/bhp077.

66. Saalmann, Y.B.; Kastner, S. Cognitive and Perceptual Functions of the Visual Thalamus, 2011. doi: 10.1016/j.neuron.2011.06.027.

67. Lázaro-Gredilla, M.; Lin, D.; Guntupalli, J.S.; George, D. Beyond imitation: Zero-shot task transfer on robots by learning concepts as cognitive programs. Science Robotics 2019, 4.

68. Jardri, R.; Denève, S. Circular inferences in schizophrenia. Brain 2013, 136, 3227–3241. doi: 10.1093/brain/awt257.

69. Jardri, R.; Duverne, S.; Litvinova, A.S.; Denève, S. Experimental evidence for circular inference in schizophrenia. Nature Communications 2017, 8. doi: 10.1038/ncomms14218.

70. Young, A.; Wimmer, R.D. Implications for the thalamic reticular nucleus in impaired attention and sleep in schizophrenia, 2017. doi: 10.1016/j.schres.2016.07.011.

71. Grossberg, S.; Hong, S. A neural model of surface perception: Lightness, anchoring, and filling-in. Spatial Vision 2006, 19, 263–321. doi: 10.1163/156856806776923399.

72. Lee, T.S.; Nguyen, M. Dynamics of subjective contour formation in the early visual cortex. Proceedings of the National Academy of Sciences of the United States of America 2001, 98, 1907–11. doi: 10.1073/pnas.031579998.

73. Lee, T.S.; Mumford, D. Hierarchical Bayesian Inference in the Visual Cortex. Journal of the Optical Society of America 2003, 2, 1434–1448.

74. Murray, S.O.; Kersten, D.; Olshausen, B.A.; Schrater, P.; Woods, D.L. Shape perception reduces activity in human primary visual cortex. PNAS 2002.

75. He, D.; Kersten, D.; Fang, F. Opposite modulation of high- and low-level visual aftereffects by perceptual grouping. Current Biology 2012, 22, 1040–1045. doi: 10.1016/j.cub.2012.04.026.

76. Kok, P.; De Lange, F.P. Shape perception simultaneously up- and downregulates neural activity in the primary visual cortex. Current Biology 2014, 24, 1531–1535. doi: 10.1016/j.cub.2014.05.042.

77. Kok, P.; Bains, L.J.; Van Mourik, T.; Norris, D.G.; De Lange, F.P. Selective activation of the deep layers of the human primary visual cortex by top-down feedback. Current Biology 2016, 26, 371–376. doi: 10.1016/j.cub.2015.12.038.

78. Bressan, P.; Spillmann, L.; Mingolla, E.; Watanabe, T. Neon Color Spreading: A Review Visual Illusions View project Self-motion perception View project Neon color spreading: a review. Perception 1997, 26, 1353–1366. doi: 10.1068/p261353.

79. Self, M.W.; van Kerkoerle, T.; Supèr, H.; Roelfsema, P.R. Distinct roles of the cortical layers of area V1 in figure-ground segregation. Current biology : CB 2013, 23, 2121–2129. doi: 10.1016/j.cub.2013.09.013.

80. Poort, J.; Self, M.W.; Van Vugt, B.; Malkki, H.; Roelfsema, P.R. Texture Segregation Causes Early Figure Enhancement and Later Ground Suppression in Areas V1 and V4 of Visual Cortex. Cerebral Cortex 2016, 26, 3964–3976. doi: 10.1093/cercor/bhw235.

81. Chen, S.; Glasauer, S.; Müller, H.J.; Conci, M. Surface filling-in and contour interpolation contribute independently to Kanizsa figure formation. Journal of Experimental Psychology: Human Perception and Performance 2018, 44, 1399–1413. doi: 10.1037/xhp0000540.

82. Johnson, J.S.; Olshausen, B.A. The recognition of partially visible natural objects in the presence and absence of their occluders. Vision research 2005, 45, 3262–3276.

83. Tang, H.; Buia, C.; Madhavan, R.; Crone, N.; Madsen, J.; Anderson, W.; Kreiman, G. Spatiotemporal Dynamics Underlying Object Completion in Human Ventral Visual Cortex. Neuron 2014, 83, 736–748. doi: 10.1016/j.neuron.2014.06.017.

84. Fyall, A.M.; El-Shamayleh, Y.; Choi, H.; Shea-Brown, E.; Pasupathy, A. Dynamic representation of partially occluded objects in primate prefrontal and visual cortex. eLife 2017, 6. doi: 10.7554/eLife.25784.

85. Ban, H.; Yamamoto, H.; Hanakawa, T.; Urayama, S.I.; Aso, T.; Fukuyama, H.; Ejima, Y. Topographic representation of an occluded object and the effects of spatiotemporal context in human early visual areas. Journal of Neuroscience 2013, 33, 16992–17007. doi: 10.1523/JNEUROSCI.1455-12.2013.

86. Hawrylycz, M.; Anastassiou, C.; Arkhipov, A.; Berg, J.; Buice, M.; Cain, N.; Gouwens, N.W.; Gratiy, S.; Iyer, R.; Lee, J.H.; Mihalas, S.; Mitelut, C.; Olsen, S.; Reid, R.C.; Teeter, C.; De Vries, S.; Waters, J.; Zeng, H.; Koch, C. Inferring cortical function in the mouse visual system through large-scale systems neuroscience. Proceedings of the National Academy of Sciences of the United States of America 2016, 113, 7337–7344. doi: 10.1073/pnas.1512901113.

87. Motta, A.; Berning, M.; Boergens, K.M.; Staffler, B.; Beining, M.; Loomba, S.; Hennig, P.; Wissler, H.; Helmstaedter, M. Dense connectomic reconstruction in layer 4 of the somatosensory cortex. Science 2019, 366. doi: 10.1126/science.aay3134.

88. Rao, R.P.; Ballard, D.H. Predictive coding in the visual cortex: A functional interpretation of some extra-classical receptive-field effects. Nature Neuroscience 1999, 2, 79–87. doi: 10.1038/4580.

89. George, D.; Hawkins, J. Towards a mathematical theory of cortical micro-circuits. PLoS computational biology 2009, 5, e1000532. doi: 10.1371/journal.pcbi.1000532.

